# Less is more! Rapid increase in plant species richness after reduced mowing of urban grasslands

**DOI:** 10.1101/805325

**Authors:** Melissa Sehrt, Oliver Bossdorf, Martin Freitag, Anna Bucharova

## Abstract

Urban lawns provide space for recreation in cities, and they are an important part of urban green infrastructures. However, most lawns are intensively managed. As only few plant species can survive the frequent mowing, urban lawns typically harbor only a limited number of plant species. To improve the biodiversity of urban lawns, it is often suggested to reduce the mowing frequency. Here, we studied the plant diversity of urban grasslands that have recently undergone management changes from mowing every few weeks to mowing only once or twice per season and compared them to intensively managed lawns. Within six years after the management changes, the grasslands with reduced mowing frequency indeed hosted 30% more plant species than intensively managed lawns, and they were more heterogeneous both within and between grasslands. Additionally, the species composition of less frequently mown grasslands shifted from common mowing-tolerant lawn species to typical meadow species. Our study thus shows that the reduction of mowing is a simple and effective tool for increasing the biodiversity in urban grasslands.

## Introduction

During the last 40 years, the movement of people towards cities greatly accelerated, and currently more than half of the global population lives in urban areas (UNFPA, 2019). Consequently, cities occupy more and more space, and urbanization has become a major threat to biodiversity worldwide (Maxwell, Fuller, Brooks, & Watson, 2016). At the same time, many people living in large cities have lost direct contact to non-urban natural landscapes, and they experience nature mainly through urban green infrastructure like parks, public lawns or private gardens. Such green urban spaces are extremely important for human well-being, and their quantity closely correlates with the quality of life in cities and the health of their inhabitants (Parker & Simpson, 2018; Tsai, Leung, McHale, Floyd, & Reich, 2019). However, not all green spaces are the same. The more biodiverse green spaces are, the larger is the positive psychological effect they provide (Southon, Jorgensen, Dunnett, Hoyle, & Evans, 2017; Wood et al., 2018; Žlender & Ward Thompson, 2017). Increasing the biodiversity of urban green spaces should thus be a win-win situation, benefitting nature and people at the same time.

A common vegetation type in urban environments are lawns – short-cut grasslands that are mown very frequently. Lawns are iconic to western cities, occupy a large proportion of urban areas and are culturally and aesthetically appreciated by most people (Ignatieva et al., 2015; Robbins & Birkenholtz, 2003). They are important spaces for recreation and outdoor activities like playing, resting, picnicking, walking or socializing (Ignatieva, Eriksson, Eriksson, Berg, & Hedblom, 2017). On the other hand, most urban lawns do not harbor much biodiversity (Aronson et al., 2017), mainly because they are too intensively managed (Klaus, 2013). As frequent mowing, fertilization and (occasional) application of pesticides are unsuitable for many grassland plants, most lawns are dominated by a limited number of tolerant species with a prostrate growth or high regrowth ability (Busch et al., 2019; Rudolph, Velbert, Schwenzfeier, Kleinebecker, & Klaus, 2017). Moreover, frequently mown plant communities tend to be rather similar across spatial scales and thus also have a low beta and gamma diversity (Chisté, Mody, Kunz, Gunczy, & Blüthgen, 2018; Gossner et al., 2016). Although such spatially homogenous lawns are considered the standard by most city managers, citizens are in fact to a certain degree open to diversification, e.g. with some urban green spaces providing space for activities and others rich sensual experiences (Ignatieva et al., 2017). Replacing part of the standard lawns by more diverse grasslands is thus a possibility for benefiting both people and nature (Klaus, 2013).

The high mowing frequency is a key factor limiting species diversity in urban grasslands (Rudolph et al., 2017). Less frequent mowing is thus a potential measure to increase plant diversity of lawns. Additional species can be sown or planted (Fischer, Lippe, Rillig, & Kowarik, 2013; Mårtensson, 2017). Alternatively, if no extra sowing or planting is done, the number of species can increase by means of natural succession, although this may take several decades, and the success of this approach depends on the availability of seeds in the seed bank and the proximity of diaspore sources in the surrounding urban landscape (Mudrák, Fajmon, Jongepierová, & Prach, 2018; Overdyck & Clarkson, 2012). Chollet et al. (2018) investigated urban grasslands after 25 years of reduced mowing frequency and natural succession, and although the diversity of grasslands significantly increased, the species richness remained rather low compared to semi-natural grasslands.

In this study, we focused on the medium-term effects of reduced mowing frequency on the plant diversity of urban grasslands in Tübingen, a city of some 80 000 inhabitants, surrounded by forests, species-rich calcareous grasslands and agricultural fields. In 2010, students of the University Tübingen established the so-called “Bunte Wiese” (Colorful Meadow) initiative that campaigns for a reduction in mowing frequency of urban grasslands to support biodiversity. Such “Colorful Meadows” are mown only once or twice per year, and small parts of the meadow are left unmown to provide shelter for overwintering insects (Unterweger, Klammer, Unger, & Betz, 2018). We compared the plant diversity of grasslands within the “Colorful Meadow” program, which experienced around six years of reduced mowing, with that of adjacent lawns mown usually 6-12 times per season. We expected that the reduced mowing frequency should (1) affect the species composition of the grasslands, (2) increase their plant species richness, (3) increase beta diversity by reducing the homogenization of vegetation both within and between grasslands, and (4) reduce species dominance and increase evenness within plant communities.

## Methods

In spring 2016, we selected 17 grasslands managed under the “Colorful Meadow” regime. Nearby each “Colorful Meadow”, we selected a grassland of similar size that was mown with the standard frequency of 6-12 times per year. Below, we refer to these two grassland types as meadows and lawns, respectively (Figure 1). All grasslands were intensively managed prior to the start of the Colorful Meadow project. The sizes of lawns and meadows were similar, on average 875 m^2^ (range 226-3376 m^2^) and 1037 m^2^ (range 259-3504 m^2^), respectively, with borders defined by natural barriers, e.g. paths, hedges, roads or buildings. In each grassland, we randomly placed five 1 m^2^ plots in which we recorded all vascular plant species and estimated their cover following the modified Braun-Blanquet scale (der Maarel, 1979). For subsequent analyses, the cover estimates were then transformed to mean percentage cover values. Using multiple small plots instead of one large plot allowed us to estimate both the heterogeneity within a grassland, based on the differences between the plots, and characterize the whole grassland based on all five plots together.

**Figure 1:**
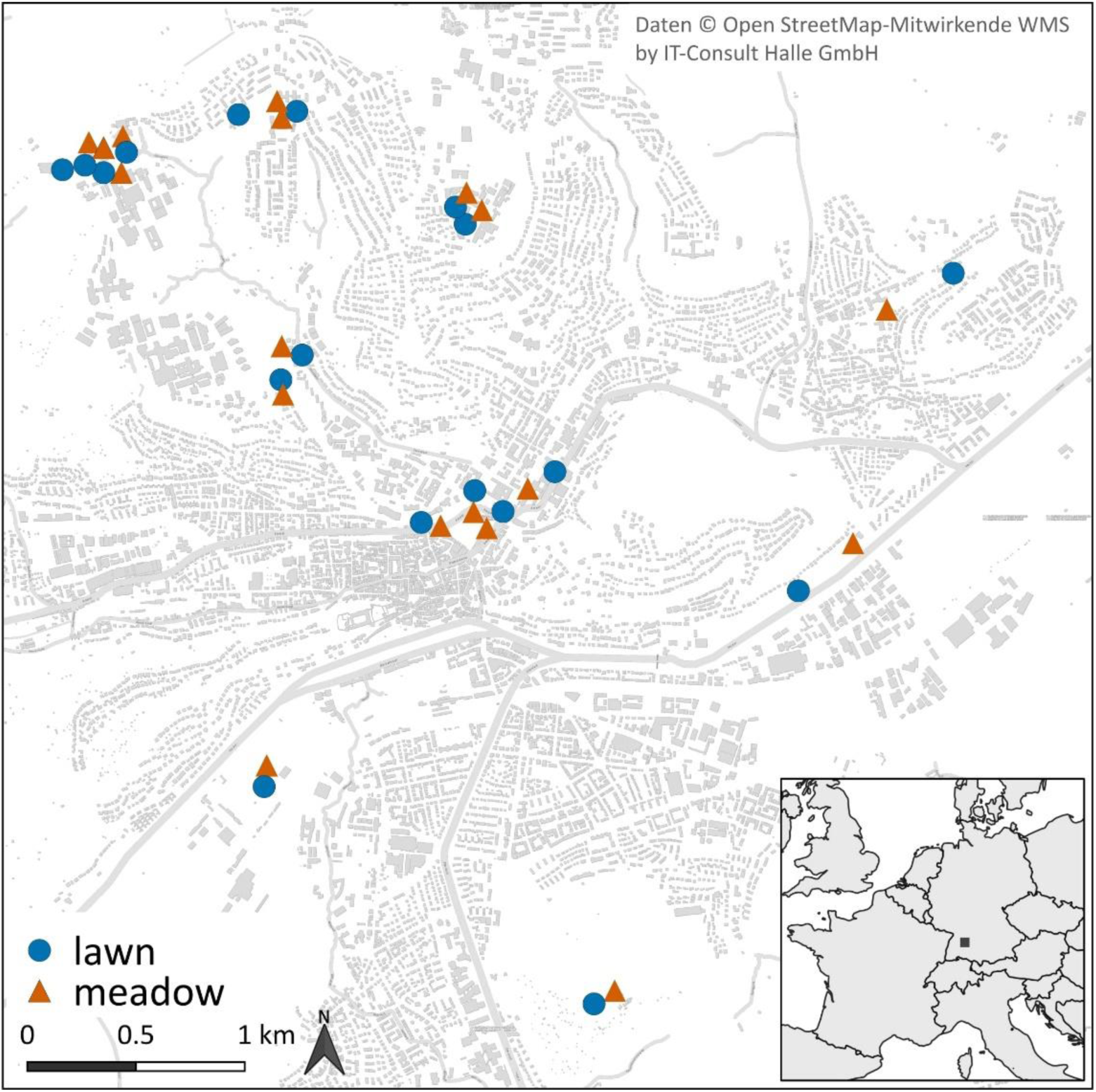
Location of lawns and meadows in the city of Tübingen.

We analyzed the data in a hierarchical fashion, from plot-level analyses of species evenness to grassland-level metrics and their relationships across all grasslands within the city. For plot-level analyses, we used species cover per plot, and for grassland-level analyses, we averaged the cover of each species across all five plots within a grassland, effectively merging the five plots into one relevé. All analyses were done in the R statistical environment (R Development Core Team, 2018) using base functions and functions from the packages *indicspecies* (Cáceres & Legendre, 2009), *lme4* (Bates, Mächler, Bolker, & Walker, 2015) and *vegan* (Oksanen et al., 2017).

First, we visualized the species composition of grasslands using detrended correspondence analysis (DCA) with rare species down-weighted. To understand which species differentiate between lawns and meadows, and are thus particularly affected by the management changes, we performed an indicator species analysis as implemented in the *indicspecies* package (Cáceres & Legendre, 2009). To obtain indicator species values, we used the square-root transformed cover values to calculate the ‘fidelity’ and ‘specificity’ of species to meadows and lawns. Fidelity is defined as the frequency of a species in a given group of grasslands, and specificity is then defined as the summed cover values of a species in a given vegetation type divided by the summed cover values of all grasslands, i.e. how exclusively a species occurs in a given vegetation type. The indicator value of a species for a given grassland type is defined as

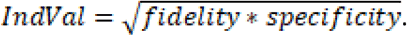

The significance of the indicator values was tested with a permutation test (N = 4999 permutations) as implemented in the *indicspecies* package.

We predicted that lawns would be species-poorer than meadows and that frequent mowing would cause homogenization both within and between grasslands. To test this, we related the number of species per grassland to the mowing regime in a linear model. Further, we calculated for each grassland the average Bray-Curtis dissimilarity between the given grassland and all other grasslands with the same mowing regime, as well as between plots within the same grassland. In the next step, we used linear models to relate these two dissimilarities to the mowing regime. All grassland-level models also included the size of the grassland as a covariate. Finally, to get some insight into vegetation structure within a plot, we calculated the evenness as the Shannon diversity index divided by the logarithm of the number of species, with high evenness indicating that the species in a community tend to have equal cover values, and low evenness indicating very unequal cover, often with dominance of few species and most others subordinate. We used linear mixed models to relate evenness to mowing regime, with mowing regime as fixed factor and, to account for plots nested within grasslands, with grassland as a random factor. All models assumed normal distribution of residuals, and we graphically evaluated that these assumptions were met.

## Results

Across all grasslands, we recorded 116 herbaceous plant species, 103 of these were found in the meadows, but only 52 in the lawns. Thirty-nine plant species occurred in both meadows and lawns, while 69 and 13 species were unique to meadows and lawns, respectively. The species composition of lawns differed from that of meadows. In the DCA, grasslands with different mowing regime were clearly separated along the first DCA axis (Figure 2).

**Figure 2:**
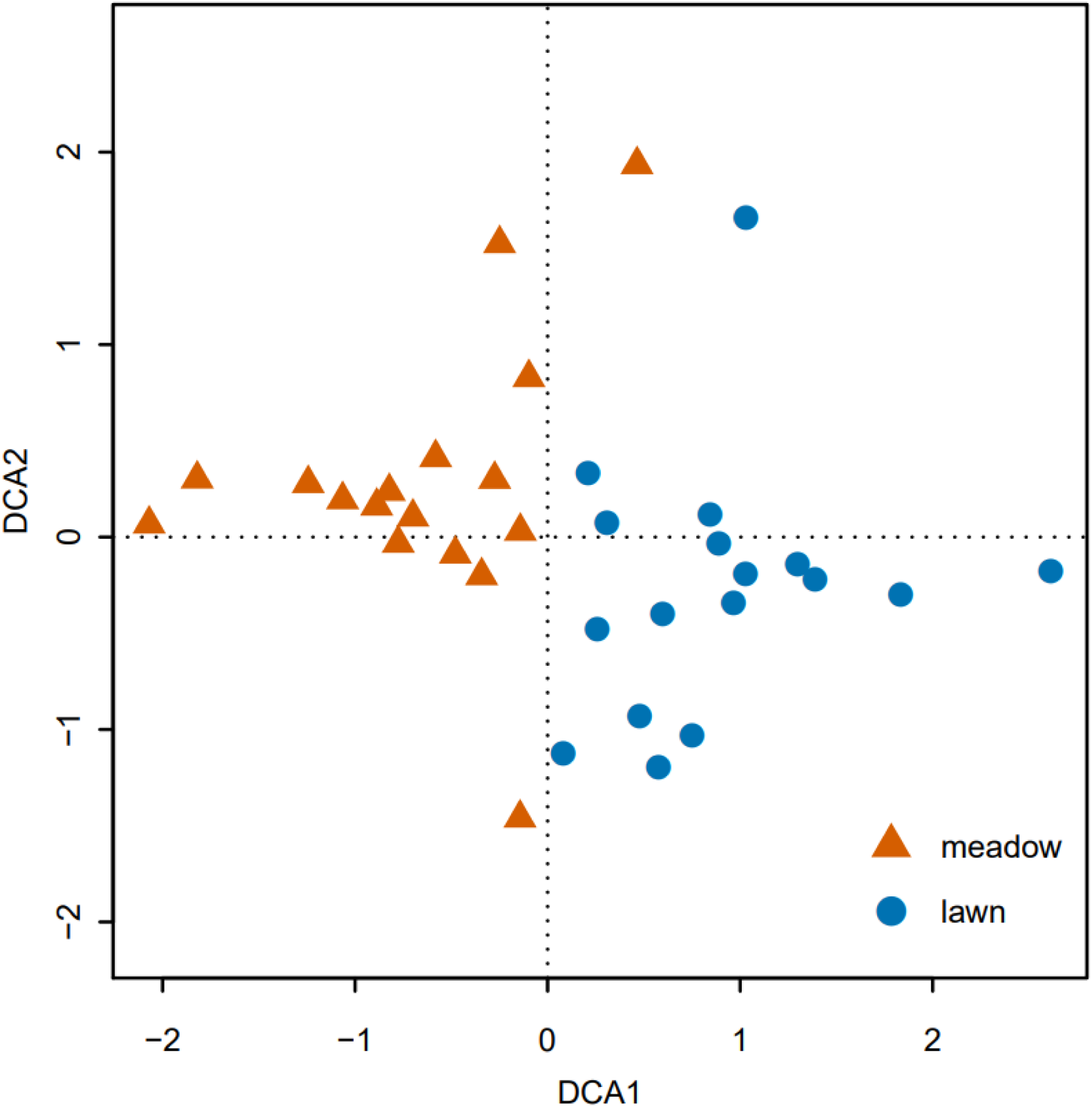
Detrended corresponded analysis visualizing differences in plant community composition across lawns and meadows. DCA1 explains 8.7 % variability, DCA2 6.8%, total inertia = 6.095.

Sixteen species were significant indicator species for either meadows or lawns. The species associated with lawns were *Bellis perennis, Lolium perenne*, *Poa annua, Prunella vulgaris* and *Trifolium repens* (Figure 3). All five species are very common and, except for *Poa annua*, occur both in lawns and meadows. On the other hand, there were 11 indicator species for meadows, of which four occurred exclusively in this grassland type: *Bromus sterilis*, *Geranium pratense*, *Rhinanthus alectorolophus* and *Trisetum flavescens* (Figure 3).

**Figure 3.**
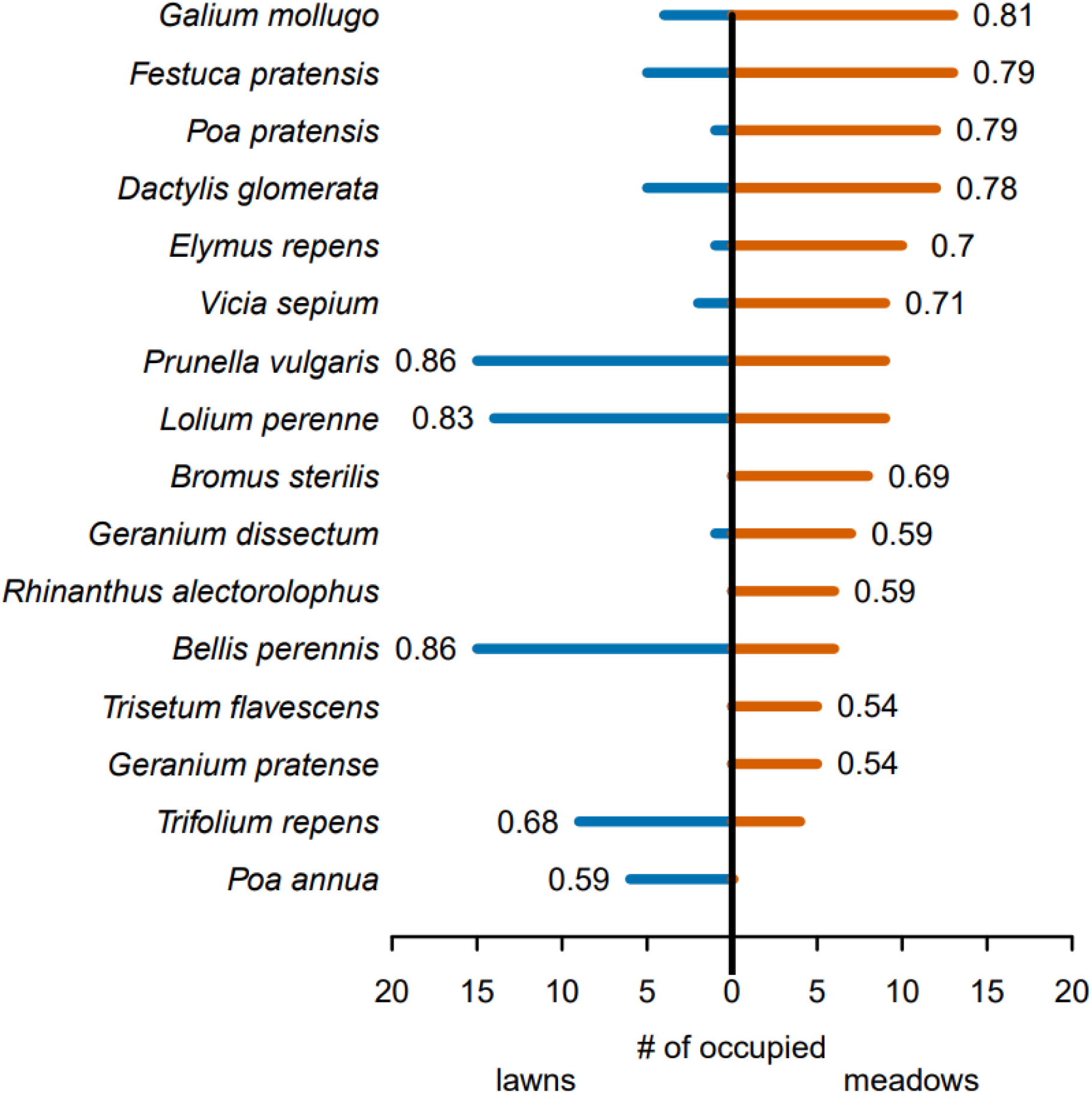
Indicator species for lawns and meadows, and frequency of their occurrence. The bars show the number of sites of a given grassland type in which species occur (out of 17 sites of each type). The numbers next to bars are indicator values of the species for either lawn or meadow. Only significant (*P*<0.05) indicator species are shown.

Species richness was higher in meadows than in lawns, with an average of 24.8 versus 17.1 species per grassland, respectively (Figure 4A, Table 1). Among grasslands within the city, lawns were more similar to each other than meadows, and also plots within one grassland were more similar to each other in lawns than in meadows (Figure 4B, C, Table 1). Finally, we found that the vegetation composition of meadows was significantly more even than that of lawns (Figure 4D, Table 1).

**Table 1:**
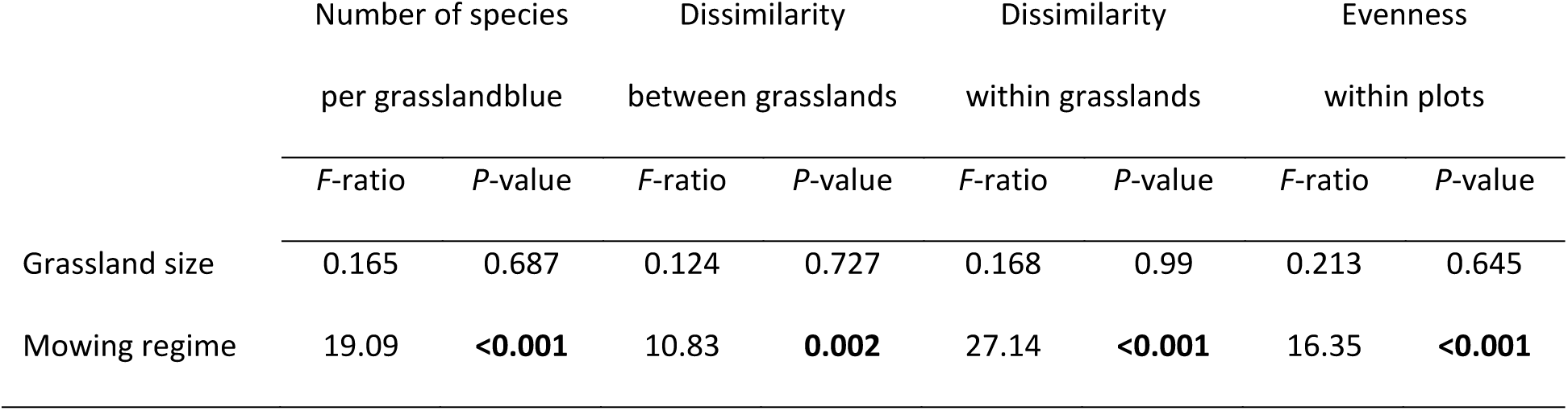
The effect of mowing regime and grassland size on the number of species per grassland, dissimilarity within and between grasslands (linear models) and species evenness within plots (linear mixed model with grassland as a random factor). Significant values (*P*<0.05) are in bold. For the analysis of evenness, the denominator degrees of freedom are 167, for all other analyses 31.

|  | Number of species<br>per grassland |  | Dissimilarity<br>between grasslands |  | Dissimilarity<br>within grasslands |  | Evenness<br>within plots |  |
| --- | --- | --- | --- | --- | --- | --- | --- | --- |
|  | *F*-ratio | *P*-value | *F*-ratio | *P*-value | *F*-ratio | *P*-value | *F*-ratio | *P*-value |
| Grassland size | 0.165 | 0.687 | 0.124 | 0.727 | 0.168 | 0.99 | 0.213 | 0.645 |
| Mowing regime | 19.09 | **<0.001** | 10.83 | **0.002** | 27.14 | **<0.001** | 16.35 | **<0.001** |

**Figure 4.**
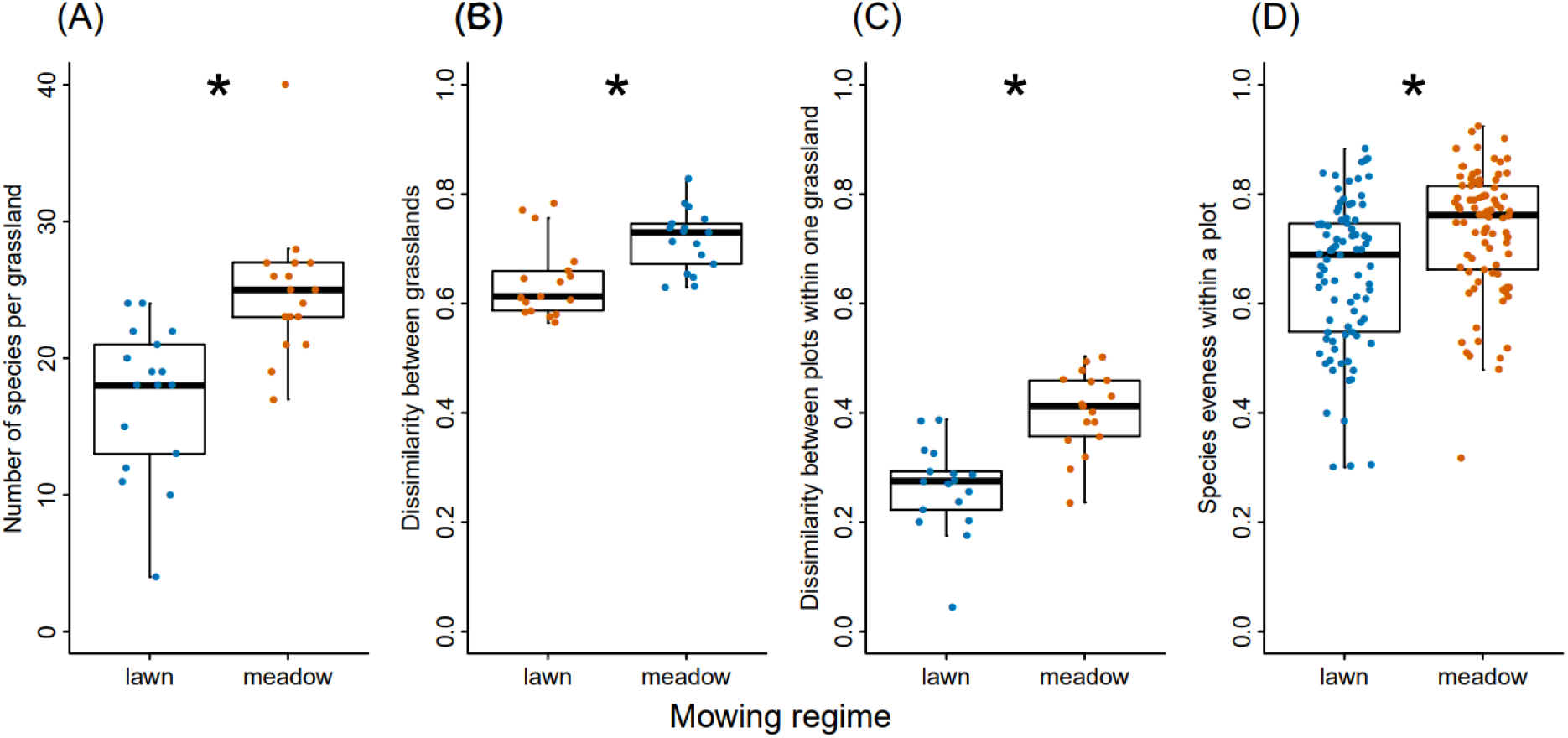
Differences in species diversity between lawns and meadows: (A) species richness, (B) dissimilarity between grasslands, (C) dissimilarity between plots within one grassland, and (D) evenness within plots. Significant differences are indicated by asterisks, see Table 1.

## Discussion

Frequent mowing is the main reason for low biodiversity of urban lawns. In this study, we have shown that the reduction of mowing frequency from every few weeks to only once or twice per season causes a species turnover and increases plant species richness of urban lawns by 30%. Moreover, the change in management also increased the spatial heterogeneity within and between grasslands. Our results are in line with previous studies showing that less intensively managed urban lawns host more plant species (Rudolph et al., 2017), and that reduced mowing alone can improve the biodiversity of urban grasslands (Chollet, Brabant, Tessier, & Jung, 2018), in our case as early as six years after the management change.

Frequent mowing restricts plant diversity in urban lawns to only few species that are able to tolerate repeated defoliation and disturbance, i.e. *Bellis perennis, Glechoma hederacea, Lolium perenne, Plantago major, Prunella vulgaris* or *Trifolium repens* (Figure 3A, Table S1). These species were present in the majority of the lawns in our study, and it is thus not surprising that lawn vegetation was rather homogenous and relatively similar to each other across the city (Figure 3B, C). Similar patterns and similar species composition have been found in urban grasslands of other European cities (Chollet et al., 2018; Politi Bertoncini, Machon, Pavoine, & Muratet, 2012; Rudolph et al., 2017). Moreover, while these species are native to Europe, many of them have been introduced to other continents where they often also dominate urban lawns, resulting in the homogenization of urban grasslands worldwide (Fischer, Rodorff, von der Lippe, & Kowarik, 2016; Wheeler et al., 2017; Zeeman, McDonnell, Kendal, & Morgan, 2017).

Reduced mowing frequency allowed more species to thrive in the urban grasslands of Tübingen. The vegetation was less dominated by the few lawn species and the species abundances were more even (see Rudolph et al., 2017 for contrasting results). Furthermore, while most lawn species were able to grow also in meadows, many additional species profited from the reduced mowing. As a result, meadows had a higher species richness, and their species pool was twice as large as that of lawn (103 vs. 52 species), a difference of similar magnitude to those reported in other urban grassland studies (Chollet et al., 2018; Rudolph et al., 2017).

Where did the additional species occurring in meadows come from? Many of them likely colonized the lawns after the reduction of the mowing frequency. Although seedling establishment is difficult in undisturbed grasslands because of microsite limitation (Klaus et al., 2017), urban grasslands are often disturbed by trampling which creates gaps necessary for seedling recruitment. New colonization was most likely the case for short-lived, ruderal species like *Bromus sterilis*, *Elymus repens*, *Galium aparine* or *Stellaria media* (Deák, Hüse, & Tóthmérész, 2016), but possibly also for some typical meadow species. Tübingen is relatively small and surrounded by species-rich meadows that are still present on steep slopes even within the city border and thus supplement the connectivity of urban green spaces (Hejkal, Buttschardt, & Klaus, 2017). These old meadows may have served as sources of diaspores for the re-colonization of the former lawns and could have contributed to the observed rapid diversity increases (Mudrák et al., 2018). Other n the other hand, many of the meadow species may have already been present in some of the grasslands already before the management change, for example *Arrhenatherum elatius*, *Trisetum flavescens* or *Geranium pratense*. Some suburbs of Tübingen have been built in areas formerly dominated by species-rich grasslands, and some urban green spaces may still be remnants of these old meadows, but now mown 6-12 times per year as urban lawns. Indeed, in several of the lawns we found typical meadows species such as *Dactylis glomerata*, *Festuca pratensis, Galium mollugo*, *Geranium dissectum*, *Poa pratenis, Ranuculus acris, Ranunculus bulbosus, Trifolium pratense* or *Vicia sepium*.

Reduced mowing frequency increased the beta diversity both at the small scale within grasslands and on the larger scale between grasslands in the city. While the effect of intensive mowing on grassland homogenization is well known (Buhk et al., 2017; Gossner et al., 2016), our results show that it is possible to revert this loss of spatial heterogeneity in plant communities by a simple and cost-effective reduction of mowing frequency (Chollet et al., 2018). The increase of beta diversity was mainly driven by species that were relatively rare in our data, growing at one or few meadows. Consequently, each meadow partly hosted a set of rather unique species (Table S1), which contrasted with the lawns that have often been dominated by the same common species.

In spite of all the positive effects of reduced mowing, the species composition of the new urban meadows still differed from the old calcareous grasslands (Mesobromion) surrounding the city. Although some typical species were found in one to three meadows, e.g. *Bromus erectus*, *Salvia pratensis*, *Thymus pulegioides* or *Pimpinella saxifraga*, many other typical species were missing (Leuschner & Ellenberg, 2018; Regierungspresidium Tübingen, 2012). As the management change was introduced only six years ago, the species composition of the meadow vegetation will further develop and presumably gain more new species in the future. Nevertheless, grasslands in the middle of a city are often associated with high level of human-mediated disturbance, and therefore urban meadows will always host novel species assemblages distinct from the reference ecosystems (Kowarik, 2011).

Although urban meadows are novel ecosystems (Klaus, 2013), this does not mean they have no conservation value. Apart of plant diversity, urban grasslands with reduced mowing frequency support a higher diversity and abundance of insects (e.g. Helden & Leather, 2004; Lerman, Contosta, Milam, & Bang, 2018; Strausz, Fiedler, Franzén, & Wiemers, 2012; Wastian, Unterweger, & Betz, 2016) and as such, they could add an important contribution to the protection of biodiversity in the Anthropocene.

## Conclusion

We have shown that reduced mowing frequency of urban lawns are apparent already within a few years after the management change. The resulting urban meadows harbor more plant species and have a less homogenous vegetation composition both within and between grasslands. While frequently-mown lawns are dominated by a few common, low-growing species, reduced management supports both ruderal species as well as tall-growing grasses and herbs characteristic for semi-natural meadows. As the species pool in cities is inevitably limited, land managers may consider sowing species from certified regional seed sources to newly created urban meadows to support regional biodiversity (Bucharova et al., 2019; Fischer et al., 2013).

## Acknowledgment

We thank Philipp Unterweger, Oliver Betz, Michael Koltzenburg and the “Bunte Wiese” initiative for their support. We are grateful to the land owners, mainly the State of Baden-Würtenberg and the city of Tübingen, which agreed to the management changes. We thank Svenja Kunze for assistance.

