## Supplementary Table 1 for "Less is more! Rapid increase in plant species richness after reduced mowing of urban grasslands"

Online supplementary material to:

Sehrt M., Bossdorf O., Freitag M. et Bucharova A.: Less is more! Rapid increase of plant species richness after reducing mowing frequency of urban grasslands. Published in Basic and Applied Ecology, 2020

**Table S1**: The list of species recorded in urban grasslands. The numbers indicate the number of lawns and meadows in which given species occurred, there was in total 17 lawns and 17 meadows in our study system.

| Species | Meadows | Lawns |
| --- | --- | --- |
| *Achillea millefolium* | 8 | 7 |
| *Agrimonia eupatoria* | 1 | 1 |
| *Agrostis* cf. *stolonifera* | 1 |  |
| *Ajuga reptans* | 7 | 7 |
| *Alliaria petiolata* | 1 |  |
| *Allium* sp. | 2 |  |
| *Alopecurus pratensis* | 2 |  |
| *Anthoxanthum odoratum* | 2 |  |
| *Arrhenatherum elatius* | 3 |  |
| *Avena sativa* | 1 |  |
| *Bellis perennis* | 6 | 15 |
| *Briza media* | 1 |  |
| *Bromus erectus* | 1 |  |
| *Bromus hordeaceus* | 1 |  |
| *Bromus sterilis* | 8 |  |
| *Campanula rotundifolia* |  | 1 |
| *Capsella bursa.pastoris* | 1 |  |
| *Cardamine pratensis* | 8 | 6 |
| *Carex caryophyllea* | 1 |  |
| *Carex flacca* | 1 |  |
| *Carex hirta* | 5 | 3 |
| *Carex muricata* | 1 | 2 |
| *Carex ovalis* | 1 | 1 |
| *Carex* sp. |  | 1 |
| *Centaurea jacea* | 1 | 1 |
| *Centaurea scabiosa* | 1 |  |
| *Cerastium holosteloides* | 6 | 8 |
| *Cirsium arvense* | 2 |  |
| *Convolvulus arvensis* | 2 |  |
| *Crepis biennis* | 3 |  |
| *Cynosurus cristatus* | 2 | 1 |
| *Dactylis glomerata* | 11 | 5 |
| *Daucus carota* | 3 |  |
| *Dipsacus fullonum* | 1 |  |
| *Elymus repens* | 1 | 1 |
| *Equisetum arvense* | 1 | 1 |
| *Eranthis hyemalis* | 1 |  |
| *Festuca pratensis* | 13 | 5 |
| *Festuca rubra* | 7 | 1 |
| *Fragaria viridis* |  | 1 |
| *Galium aparine* | 4 |  |
| *Galium mollugo* | 13 | 4 |
| *Galium vernum* | 2 | 1 |
| *Geranium pyrenaicum* | 1 |  |
| *Geranium dissectum* | 7 | 1 |
| *Geranium pratense* | 5 |  |
| *Geum urbanum* | 7 | 3 |
| *Glechoma hederacea* | 13 | 15 |
| *Hedera helix* | 1 | 2 |
| *Helictotrichon pubenscens* | 1 |  |
| *Heracleum sphondylium* | 1 |  |
| *Hieracium spec.* | 1 |  |
| *Holcus lanatus* | 4 |  |
| *Hypericum perforatum* | 1 |  |
| *Hypochaeris radicata* | 1 | 1 |
| *Knautia arvensis* | 2 | 1 |
| *Lamium purpurea* | 1 |  |
| *Latyrus pratensis* | 2 |  |
| *Leontodon autumnale* | 1 | 1 |
| *Leontodon hispidus* | 1 | 1 |
| *Leucanthemum vulgare* | 5 | 4 |
| *Listera ovata* | 1 |  |
| *Lolium perenne* | 9 | 14 |
| *Lotus corniculatus* | 6 | 5 |
| *Luzula campestris* | 1 |  |
| *Lysimachia nummularia* | 8 | 4 |
| *Malva moschata* | 1 |  |
| *Medicago lupolina* | 5 | 4 |
| *Myosotis arvensis* | 1 |  |
| *Ononis spinosa* |  | 1 |
| *Origanum vulgare* | 1 |  |
| *Oxalis acetosella* | 1 |  |
| *Pilosella sp.* |  | 4 |
| *Pimpinella saxifraga* | 1 |  |
| *Planatgo media* |  | 1 |
| *Plantago lanceolata* | 15 | 12 |
| *Plantago major* | 3 | 7 |
| *Poa annua* |  | 6 |
| *Poa palustris* | 6 | 5 |
| *Poa pratensis* | 12 | 1 |
| *Poa trivialis* | 7 | 2 |
| *Potentilla anserina* | 2 |  |
| *Potentilla reptans* | 12 | 9 |
| *Prunella vulgaris* | 9 | 15 |
| *Ranunculus acris* | 11 | 8 |
| *Ranunculus bulbosus* | 6 | 5 |
| *Ranunculus ficaria* | 3 | 1 |
| *Rhinanthus alectorolophus* | 6 |  |
| *Rumex acetosa* | 2 | 3 |
| *Rumex obtusifolius* | 1 |  |
| *Rumex crispus* | 1 | 2 |
| *Salvia pratensis* | 2 | 1 |
| *Senecio viscosus* |  | 2 |
| *Silene latifolia* | 1 |  |
| *Sonchus oleraceus* | 1 |  |
| *Stellaria media* | 2 |  |
| *Tanacetum corymbosum* | 1 | 1 |
| *Tanacetum vulgare* | 1 |  |
| *Taraxacum sect.* | 11 | 13 |
| *Thlapsi arvense* | 1 |  |
| *Thymus pulegiodes* | 1 | 1 |
| *Trifolium dubium* | 3 | 4 |
| *Trifolium pratense* | 12 | 11 |
| *Trifolium repens* | 3 | 9 |
| *Trisetum flavescens* | 5 |  |
| *Veronica arvensis* | 3 | 1 |
| *Veronica chamaedrys* | 9 | 6 |
| *Veronica filiformis* | 2 | 4 |
| *Veronica hederifolia* | 4 | 2 |
| *Veronica persica* | 6 | 2 |
| *Veronica serpyllifolia* | 1 | 2 |
| *Vicia cracca* | 1 |  |
| *Vicia sepium* | 9 | 2 |
| *Vicia tetraspermum* | 1 |  |
| *Viola arvensis* | 1 |  |
| *Viola* sp. | 2 | 7 |
